# Transfer learning of deep neural network representations for fMRI decoding

**DOI:** 10.1101/535377

**Authors:** Michele Svanera, Mattia Savardi, Sergio Benini, Alberto Signoroni, Gal Raz, Talma Hendler, Lars Muckli, Rainer Goebel, Giancarlo Valente

## Abstract

**Background:** Deep neural networks have revolutionised machine learning, with unparalleled performance in object classification. However, in brain imaging (e.g. fMRI), the direct application of Convolutional Neural Networks (CNN) to decoding subject states or perception from imaging data seems impractical given the scarcity of available data.

**New method:** In this work we propose a robust method to transfer information from deep learning (DL) features to brain fMRI data with the goal of decoding. By adopting Reduced Rank Regression with Ridge Regularisation we establish a multivariate link between imaging data and the fully connected layer (`fc7`) of a CNN. We exploit the reconstructed `fc7` features by performing an object image classification task on two datasets: one of the largest fMRI databases, taken from different scanners from more than two hundred subjects watching different movie clips, and another with fMRI data taken while watching static images,

**Results:** The `fc7` features could be significantly reconstructed from the imaging data, and led to significant decoding performance.

**Comparison with existing methods:** The decoding based on reconstructed `fc7` outperformed the decoding based on imaging data alone.

**Conclusion:** In this work we show how to improve fMRI-based decoding benefiting from the mapping between functional data and CNN features. The potential advantage of the proposed method is twofold: the extraction of stimuli representations by means of an automatic procedure (unsupervised) and the embedding of high-dimensional neuroimaging data onto a space designed for visual object discrimination, leading to a more manageable space from dimensionality point of view.

## 1. Introduction

A long-standing goal of cognitive neuroscience is to unravel the brain mechanisms associated with sensory perception. Cognitive neuroscientists often conduct empirical research using non-invasive imaging techniques, among which functional Magnetic Resonance Imaging (fMRI) or Electroencephalography (EEG), to validate computational theories and models by relating sensory experiences, like watching images and videos, to the observed brain activity. Establishing such relationship is not trivial, due to our partial understanding of the neural mechanisms involved, the limited view offered by current imaging techniques, and the high dimensions in both imaging and sensorial spaces.

A large amount of statistical approaches have been proposed in the literature to accomplish this task; in particular, in the last two decades great attention has been given to *generative* (also referred to as *encoding*) and *discriminative* (*decoding*) models, that have different aims, strengths and limitations (see [1]). Encoding models aim at characterising single units response harnessing the richness of the stimulus representation in a suitable space, and can thus be used to model the brain response to new stimuli, provided that a suitable decomposition is available. On the other hand, decoding models solve a “simpler” problem of discriminating between specific stimulus types and are better suited, when the available training data are relatively scarce, in capturing correlations and interactions between different measurements and are thus optimised for prediction (see [2], [3]).

In both approaches there is a heavy emphasis on ideas and algorithms developed in machine learning (ML). This field has enormously benefited from the recent development of Deep Neural Networks (DNN), originally designed to tackle object classification tasks. By integrating a series of differentiable layers, these networks exploit multi-level feature extraction (from low level e.g., color and texture, to higher level features, more category oriented) becoming an end-to-end, often defined as “biologically inspired”, classification tool. Historically, the deep learning community has always been inspired by the brain mechanisms while developing new methods and cognitive neuroscience can provide validation of AI techniques that already exist. The two communities therefore share now many common research questions [4, 5, 6]: for example, how the brain transforms the low-level information (colors, shapes, etc.) into a certain semantic concept (person, car, etc.) is an important research topic for both.

When dealing with visual stimuli, in the last few years the brain imaging community has been making more and more use of deep neural networks. To this avail, several studies attempted to relate these models with brain imaging data revealing interesting similarities between DNN architectures and the hierarchy of biological vision [7]. An interesting study showed how a DNN resembles representational similarity of Inferior Temporal (IT) intra- and inter-categories [8]. Another relevant study [9] described how a CNN captured the stages of human visual processing in time and space from early visual areas towards the dorsal and ventral streams.

Alongside the research that investigates the computations performed in the visual pathway by comparing the behaviour of deep neural networks and measured neural responses, another active area of research focuses more on examining how far these methods can be applied to brain imaging to improve existing statistical approaches. In this respect, most of the applications can be found in the context of *encoding* models, where each training stimulus is described using an {*m*}-dimensional representation and a generative model based on such representation is estimated at each brain location. Representing the stimuli with more abstract features, derived from deep neural networks, the authors in [10] achieved better performance in reconstructing brain activity, using the dataset of [11], where Gabor pyramid wavelets were used to decompose visual stimuli. Similarly, DNN-derived features have been used in [12], which introduced new classes of encoding models that can predict human brain activity directly from low-level visual input (*i.e.*, pixels) with ConvNet [13]. In [14] encoding models were developed to predict fMRI single-voxel response, extending *de facto* [10] to movie viewing, trying to capture the dynamic representations at multiple levels. In [15] authors presented an encoding model by which, starting by Convolutional Neural Network (CNN) layer activations and using ridge regression with linear kernel, they predict BOLD fMRI response, employing two different databases ([11] and [16]). In [17] the authors presented a novel image reconstruction method, in which the pixel values of an image are optimised to make its CNN features similar to those decoded from human brain activity at multiple layers. A further example of encoding came from [18], in which the prediction of brain response is done multi-subject and using Bayesian incremental learning.

Whereas *encoding* models have greatly benefited from the inclusion of DNN-derived features in the modeling pipeline, *decoding* models have not yet exploited the full potential offered by them. Despite the fact that DNN are discriminative models, there is an obvious reason why they have not been extensively used in decoding applications: the number of samples typically available in the imaging studies is far too low to be able to successfully train a deep network. Even when pooling together multiple sites, a deep neural network does not outperform a much simpler kernel ridge regression with L2 regularisation [19]. An early study that exploits the idea of using CNN representations in decoding is [14], in which convolutional (conv) and fully connected (`fc`) layers are compressed before performing prediction and subsequently classification, with good within-subject performance.

In this work we propose an approach in which the richness of feature representation provided by deep artificial neural networks can be harnessed to enhance the performance of fMRI-based decoding. Since the sheer amount of samples needed to train a deep neural network is simply not available in imaging experiments, we propose instead to use a CNN to extract different visual data content representations and subsequently link these representations with brain data, thus performing an {*n*} to {*m*} mapping (*n* = voxels, *m* = visual features), followed by prediction on new data. Importantly, we implement a simple but effective method that involves the prediction, rather than the stimulus itself, of an intermediate representation of the stimulus in order to partially transfer information, or simply a property, from its representation to the initial data. This approach, well know in Deep Learning as *transfer learning*, has the ability to allow an abstraction from the raw data, potentially expanding the analysis also to unseen data, since stimuli representations may more easily address unsupervised learning tasks [20, 21, 22]. We therefore build on the intuition from [14], doing an across subjects prediction, comparing different multivariate and multiple linking methods, optimising the hyper-parameters of the linking, and using voxels from all brain without *a priori* selecting areas of interest.

To transfer information from the DNN features to imaging data several approaches are available, most of which are based on ideas of dimensionality reduction and latent structures. Very common examples in multimodal neurophysiological data are provided by Canonical Correlation Analysis (CCA) [23] and Partial Least Square (PLS) [24], which project the original data sets in new spaces, emphasising, respectively, the role of correlations and covariance among the projected data. Additional methods, like Independent Component Analysis (ICA) [25, 26, 27] or Dictionary Learning/Sparse coding [28, 29], try to identify the set of source signals which produce the set of mixed signals read in measurements.

By transferring information from CNN to imaging data, we show that it is possible to achieve better discrimination, as compared with using imaging data alone. To demonstrate the validity of the proposed approach we make use of two different datasets. The first, from [3], involves free viewing of movie excerpts and is characterised by a large number of subjects. On the second dataset, based on static images presentation [11] we instead implement within-subject prediction, performing decoding of visual categories.

## 2. Materials and Methods

The general idea behind the proposed approach is presented in Figure 1. To create a training set we analyse the images (or movie data) by means of a CNN architecture and extract deep features from the last fully-connected layer (from now on, identified as `fc7`). Since we are interested in performing decoding and classifying visual object classes, we select `fc7`, the penultimate CNN layer before classification, which is considered as a highly representative feature of the object class and shape [30]. The objective is to robustly learn, by a linking method, an association between these two high dimensional datasets; this link enables us to predict the last fully-connected layer of a CNN (`fĉ7`) using brain data from fMRI of untested subjects watching unseen images (or movies).

**Figure 1:**
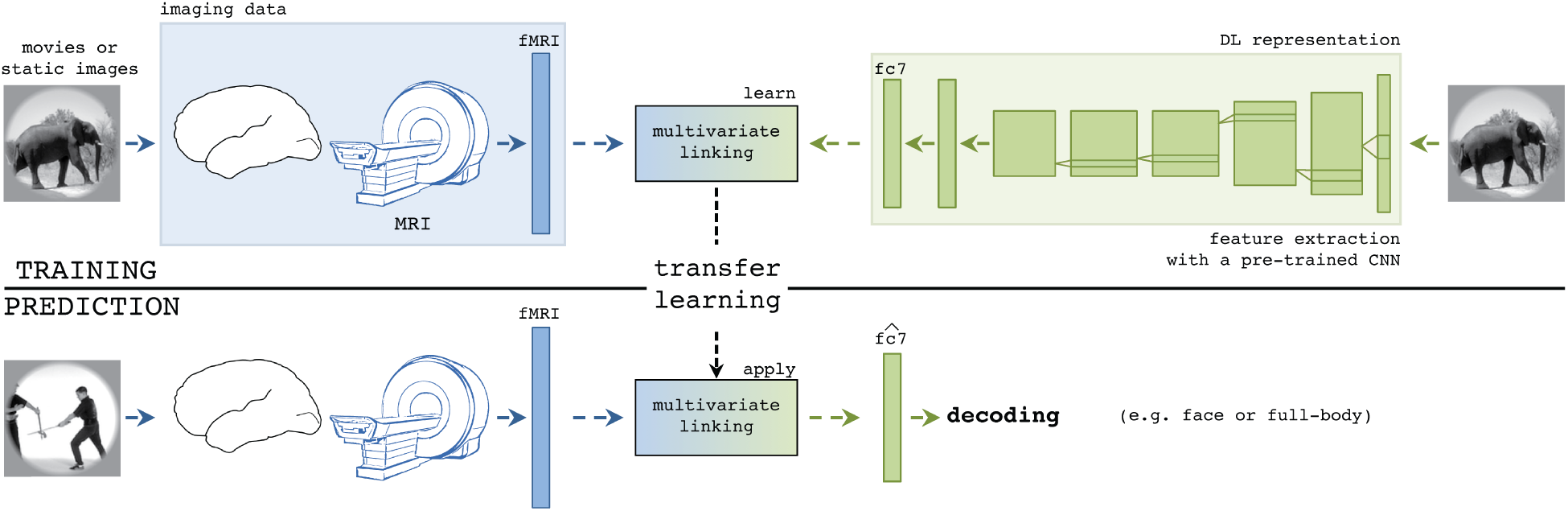
Framework description for mapping fMRI to and from `fc7` deep features. Deep learning features are extracted from a pre-trained CNN and image data are collected using fMRI. The training phase learns the ability to reconstruct `fc7` from brain data through multivariate linking.

We validated the approach on two fMRI datasets: an image dataset widely used in the context of visual categorisation, encoding and DNN modeling [11], and a movie dataset [3], respectively in Section 2.1.2 and 2.1.1. The rationale behind using a movie viewing dataset is that, whereas most of the current imaging studies use strictly controlled conditions as stimuli employing single images surrounded by controlled contours and interleaved with rest period, the natural everyday experience of human beings is closer to videos than images. Therefore, the neural responses elicited by watching a more ecologically valid stimulus, such as a movie, are more representative of normal functioning of the brain. We thus test the presented approach in this very challenging scenario, using one of the vastest database of natural movies ever used so far in the context of fMRI decoding [3], with ∼ 37, 500 time points, without imposing a priori selection of brain regions (*i.e.*, ∼42, 000 voxels), and using, in the test phase, novel movies and unseen subjects.

To perform the linking, different high-dimension multivariate regression methods are tested and compared in Section 2.2. In this section, we furthermore illustrate how to tune the hyperparameters of the model, which is particularly challenging in the movie dataset, given the large amount of time points. Finally, two example of classification, based on the transfer learning approach developed in this work, are shown in Section 2.3.

### 2.1. fMRI datasets

#### 2.1.1. Movie Dataset

##### Imaging data description

We use a set of stimuli consisting of 12 film clips between 5 – 10 minutes in duration, for a total length of ∼ 72 minutes. Movie data are part of a larger dataset collected for projects examining hypotheses unrelated to this study. All clips adhere to the so-called classical Hollywood-style of film making, characterised by continuity editing, the use of abundant emotional cues, and an emphasis on narrative clarity. In Table 1 relevant information about movies and subjects are reported: title, duration and few subject properties. For subject clustering, acquisition details, and pre-processing steps, please refer to original works in [31, 3].

**Table 1:**
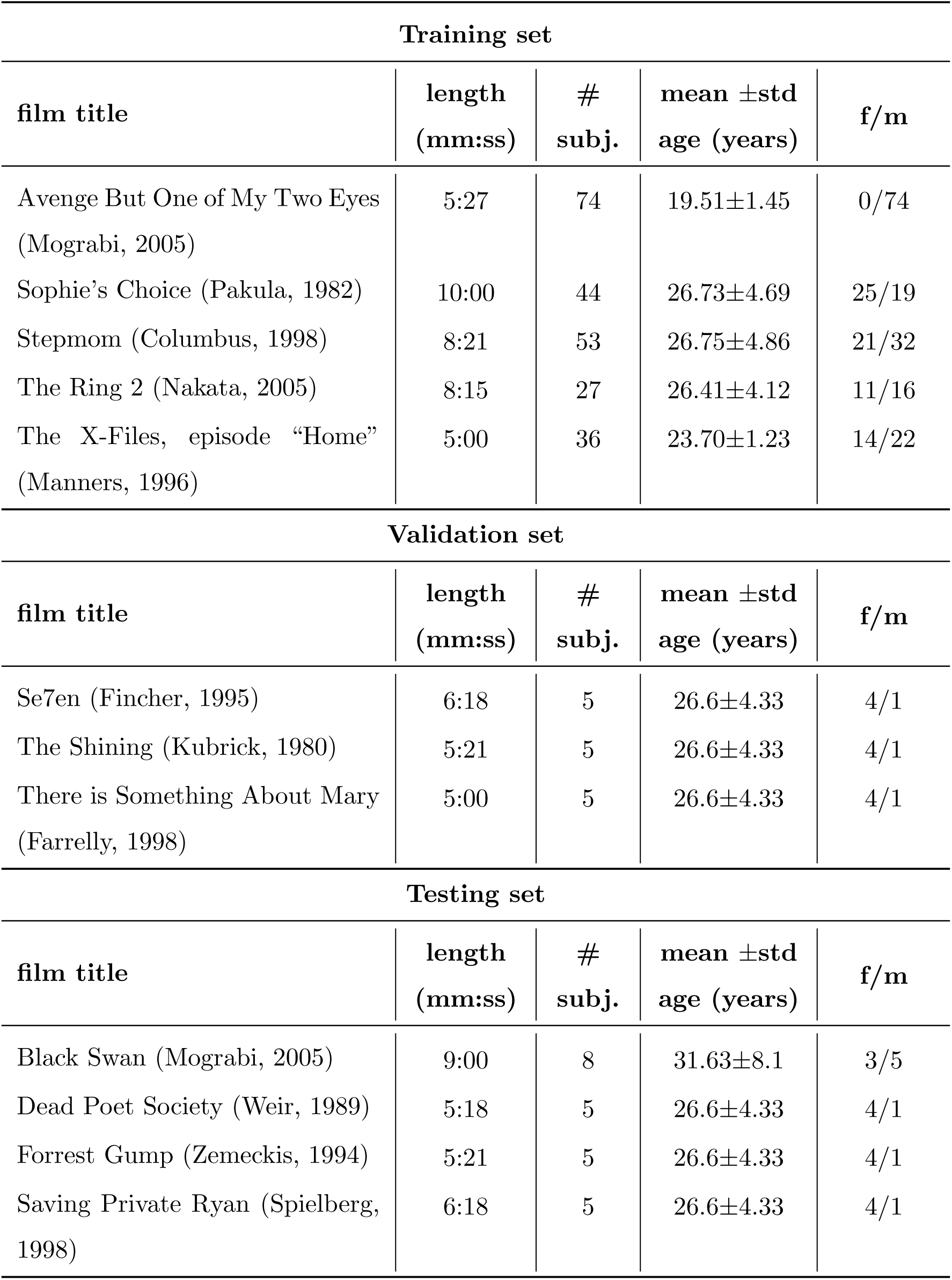
Movie dataset: details on the movie material and samples used in the study.

fMRI data are collected from several independent samples of healthy volunteers with at least 12 years of education using a 3 Tesla GE Signa Excite scanner. Due to technical problems and exaggerated head motions (1.5 mm and 1.5° from the reference point) only stable data are included. Functional whole-brain scans were performed in interleaved order with a T2*-weighted gradient echo planar imaging pulse sequence (time repetition [TR]/TE = 3, 000/35 ms, flip angle = 90, pixel size = 1.56 mm, FOV = 200 × 200 mm, slice thickness = 3 mm, 39 slices per volume). Data are pre-processed and registered to standardised anatomical images via Brainvoyager QX version 2.4 (Brain Innovations, Maastricht, Netherlands). Data are high pass filtered at 0.008 Hz and spatially smoothed with a 6 mm FWHM kernel. We confined the analysis using a gray matter mask based on an ICBM 452 probability map [32] thresholded to exclude voxels with probability lower than 80% of being classified as gray matter (thus encompassing both cortical and brain stem regions) obtaining a fMRI data with *∼* 42, 000 voxels.

##### CNN Feature extraction

Nowadays, many applications in computer vision use CNNs for feature extraction: passing the image through a network, reading some activations, and using them to represent the image or feeding the features to a classifier. The choice on which layer to extract depends on the task under examination: convolutional layers act by creating a bank of filters which return shift-invariance features, exploiting the intrinsic structure of images; fully connected layers learn a representation closer to categorical visual classes. Since we are interested in performing decoding and classifying visual object classes, we select `fc7`, the penultimate CNN layer before classification. The features are extracted after ReLu, *i.e.*, thresholded, thus obtaining a sparse representation of the object class, even if a comparison with and without rectified linear unit layer (ReLu) in done in Section 3.2. The entire framework here proposed is expandable to different layers without changing the structure of the methods.

Features are extracted and collected from video frames as described in Figures 2. First, each processed frame feeds a *faster R-CNN* network ([33]). Multiple objects, together with their related confidence values and last fully connected layer (`fc7`), are therefore extracted from each processed frame at different scales and aspect ratios. Since it is possible to have in one frame multiple detections of the same object class (as in Figure 2 for the class “person”), for each class only the `fc7` layer of the object with maximum confidence is kept. For this work only “person” class is considered, obtaining a 4, 096 dimension feature vector from each frame.

**Figure 2:**
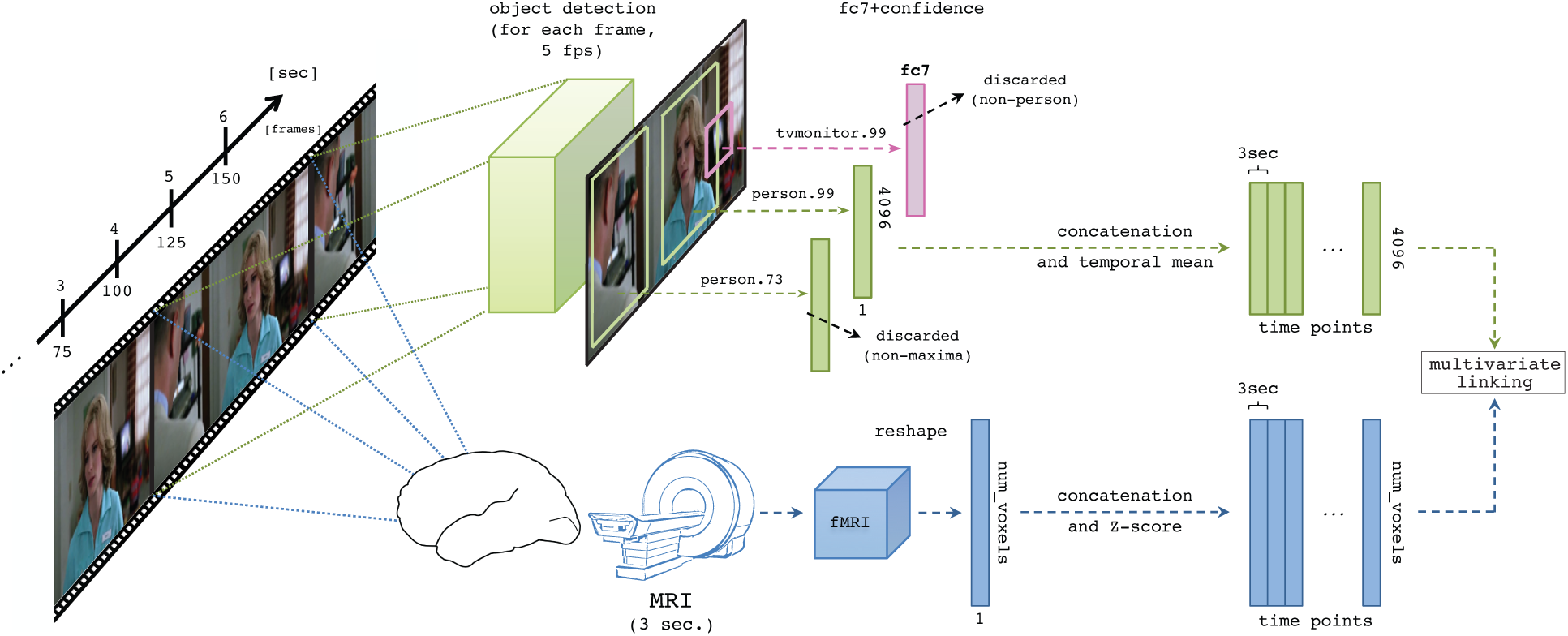
Framework description for mapping fMRI to and from `fc7` deep features, thus enabling decoding and encoding, respectively. Video features are extracted for each processed frame in the video (5fps) and temporally averaged (3s) in order to be aligned with voxel time courses.

The whole procedure is performed at a frame rate of 5*fps* on every movie clip. As shown in Figure 2, in order to properly align the `fc7` feature matrix with the fMRI data resolution (3*s*), `fc7` feature vectors are averaged on sets of 15 frames. Different subjects and different movies are concatenated in time dimension, keeping valid the correspondence between fMRI and visual stimuli: subjects watching equal movie share the same `fc7` features but different fMRI data.

In this work, we assume that subjects are only focusing on persons in the scene; assuming that the attention of the subjects while watching movies is directed to the classes in analysis is an assumption which is corroborated by many studies in literature. In fact, in cinema studies human figures are well known to be central to modern cinematography [34], especially in Hollywood movies, and are often displayed in the center of the frame [35]. Moreover, in brain imaging, the work in [36] showed that the correlations between subjects watching the same movie are very similar not only in eye movements, but also in brain activities, suggesting similar focus of attention across participants. It is important to stress that, even if we focus on person class only with the movie dataset, the proposed work can be expanded to different classes for different experiments without changes in the framework architecture.

#### 2.1.2. Static images dataset

##### Imaging data description

In order to test the generality of the method in a more common and controlled situation, we challenge the proposed model also on static images. In [11], Kay and colleagues introduced one of the first successful encoding method applied to images. In the original work, a model based on Gabor pyramid wavelets was trained to predict every voxel response separately. The entire database includes 1, 750 training and 120 validation images.

Along with the publication and images, authors made available also the estimated fitted General Linear Model (GLM) betas per voxel. The provided responses for each voxel have been z-scored, so for a given voxel the units of each “response” are standard deviations from that voxel’s mean response. Around 25, 000 voxels in or near the cortex were selected for each of the two subjects. Different works have made use of this database, for instance see [37], or [10].

The experimental design, MRI acquisition protocol, and preprocessing of the data are identical to those described in these studies. The study collected fMRI data for two male subjects (*S*_1_ and *S*_2_), watching selected training and testing images. Data were acquired using a 4 T INOVA MR scanner and a quadrature transmit/receive surface coil. Eighteen coronal slices were acquired covering occipital cortex (slice thickness 2.25 mm, slice gap 0.25 mm, field of view 128 × 128 mm^2^). fMRI data were acquired using a gradient-echo EPI pulse sequence (matrix size 64 × 64, TR 1 s, TE 28 ms, flip angle 20°, spatial resolution 2 × 2 × 2.5 mm^3^). See [11] for details of BOLD response estimation, voxel selection, and ROI definition.

Despite only two subjects are available, limiting the results generality across subjects, the outcome is still informative for our work, since the database is composed by many images.

##### CNN Feature extraction

We extract two sets of features from image material. The first, following the same procedure used for movie clips, involves faster R-CNN, and results in a representation of the “person” class, with the final goal of performing classification (results reported in Sec. 2.3). The second set of features is instead obtained by another CNN. A common choice for a classification task is nowadays to use VGG-16 [38], which has been pre-trained on ImageNet database [39]. To prove the association ability of the method between deep features and brain data, we choose to extract a general image description using this network. Originally Kay’s database does not come with a ground-truth containing annotations on video object classes. Therefore to provide a valid ground truth for the “person” class, three different human annotators created annotations which were then mediated, for the classification “person” vs “no-person”. For other visual object classes, such as those present in ImageNet database, the classes present in images were heavily unbalanced in cardinality, making the classification unreliable. To validate the reconstruction performance, we report correlation result in Section 3.3.

### 2.2. Linking methods

The association between the fMRI data and the deep features `fc7` (see multivariate linking box in Figure 1) can be learnt using multivariate linking methods. Canonical Correlation Analysis (CCA) [23] is often used in this respect [40, 41, 42, 43, 44], as it allows projecting one dataset onto another by means of linear mapping, which can be further used for categorical discrimination and brain model interpretations. CCA aims at transforming the original datasets by linearly projecting them onto new orthogonal matrices whose columns are maximally correlated. To capture nonlinear relationships between data, or to simply make the problem more tractable, kernel versions are often used, which consist in projecting (linearly or non-linearly) data onto a different space before performing CCA. In addition, regularised versions of CCA allow to extend the method when the number of dimensions is close to or exceeds the available time points. In this work we used the implementation proposed in [45].

Similarly, Partial Least Square (PLS) [24] maximises the covariance of the matrices in the new spaces and different extensions of the method are particularly suited to the analysis of relationships between measures of brain activity and of behaviour or experimental design [46].

Among other high-dimensional approaches, multivariate linear regression ({*n*} to {*m*}) is a widely employed strategy. Multivariate linear regression is the extension of the classical multiple regression model to the case of both multiple (*m* ≥ 1) responses and multiple (*n* ≥ 1) predictors (in this case *n* = number of voxels (∼42, 000), *m* = size of `fc7` (4, 096)). Among all approaches, a promising and elegant formulation can be found in the work of [47], with a reduced `rank` ridge (RRRR) approach for multivariate linear regression and it is particularly suited for the current problem. Starting from the assumption that the response matrix is often intrinsically of lower rank, due to the correlation structure among the prediction variables, the method combines an L2 norm penalty (*i.e.*, ridge) with the reduced rank constraint on the coefficient matrix, efficiently handling the high-dimensional problem we face. For a complete formulation of RRRR and the related mathematical proof see [47] (in Appendix A a short mathematical formulation is provided).

In this work we compared Canonical Correlation Analysis (CCA) with different kernels (linear, gaussian, and polynomial), Partial Least Square (PLS) and Reduced `Rank` Ridge approach for multivariate linear Regression (RRRR). Short descriptions of these methods, together with references and toolboxes are reported in Table 2. For an extensive description of these and other methods, and their use on brain data, please refer to [48].

**Table 2:** Multivariate mapping methods: state of the art in [48].

| Method | motivation | optimisation criteria | limitations | reference | library |
| --- | --- | --- | --- | --- | --- |
| CCA | Determines correlated sources across two data sets without considering variation information. | The original datasets are linearly projected, using two matrices, onto new orthogonal matrices $U$ and $V$ whose columns are maximally *correlated*. | CCA may not be appropriate when $n$, $m \approx t$ or $n$, $m \gg t$ (solved using regularisation and kernel versions). Constrains the demixing matrix to be orthogonal, hence limiting the search space for the optimal solution. | [23] | [45] |
| RRRR | Selects latent variables that explain as much response ($Y$) variation as possible. Rank constraint encourages dimension reduction. Ridge penalty ensures that the estimation of $B$ is well-behaved even with multicollinearity. | Minimises sum of square errors with rank constraint on $B$ (shrinking) and with L2 norm penalisation on regression coefficients (ridge penalty), maximising the correlation. | Ignores the predictors for the purposes of factor extraction. With respect to CCA it is less interpretable and posits an intrinsic directionality in the relationship between datasets. | [47] | n.a. |
| PLS | Determines correlated sources across two data sets considering variation information. Used also to determine which part of the observations are related directly to another set of data. | Maximises the *covariance* between corresponding sources across two data sets. Finds a linear regression model by projecting the predicted variables and the observed variables to a new space. | Higher covariance between two corresponding latent variables does not necessarily imply strong correlations between them. Little is known about effective means of penalization to ensure sparse solutions and avoid overfitting. | [24] | [49] |

#### 2.2.1 Hyper-parameters optimisation

Establishing the link between fMRI data and fc7 features involves the choice of many hyper-parameters, that can be optimised. Noteworthy, we here use the term model “hyper-parameters”, with respect to simply model “parameters”, to distinguish those values that cannot be learnt during training, but are set beforehand *e.g.*, the regularisation terms or the number of hidden components. Whereas the use of the very large movie dataset (2.1.1) makes it possible to refine and optimise on the training data these hyper-parameters, the large amount of available data makes it computationally unfeasible to use grid-search or random-search approaches. The solution here adopted makes use of a highly efficient sequential optimisation technique based on decision trees taken from [50].

This approach provides a faster and more cost-effective optimiser by exploiting the underlying hyper-parameter space by means of decision trees; this allows to describe the relation of the target algorithm performance with respect to the hyper-parameters, thereby finding the minimum with as few evaluations as possible. In practice, several random points are extracted from the parameter probability distributions and several models are trained (on the training set); after performance evaluation (on the validation set), the decision trees model computes the next best point, minimising the cost function. In our case, we use the optimiser to maximise the mean correlation between original `fc7` and reconstructed `fĉ7` across all validation movies (described in Table 1).

In the case of CCA with different kernel versions with ridge regularisation we used the package provided in [45]. Despite the large amount of available computer memory (256GB), we could only use half of the time points of the training set (one point every two), since the CCA - only - method requires a large memory and a long time to be trained. The CCA hyper-parameters to optimise are: the regularisation term, the number of components, and - in case of kernels - its degree, for the polynomial kernel, or sigma, for the gaussian kernel.

The PLS regression model is trained using the code in [49], and by optimising the number of components, whereas for the RRRR, implemented in Python^1^ based on the R code provided by Mukherjee et al. [47], the hyper-parameters optimised are the rank and the L2 regularisation weight. After this comparison was conducted, we addionally performed a more in-depth hyperparameter optimisation for the RRRR algorithm; the optimised hyperparameters are described in Table 3), together with their range.

**Table 3:** Model training: hyper-parameter selection. List and description of all the hyper-parameters to be optimised during training and list of related figures.

| Parameter | Description | Range | Figure |
| --- | --- | --- | --- |
| `rank` | Rank of the regressor | 1 – 4096 | 4-(a) |
| `reg` | Regularisation term | $10^{-12: +12}$ | 4-(a) |
| `time_shift` | Temporal alignment (in volumes) between video features and fMRI samples: a negative value means that `fc7` are delayed with respect to fMRI time series | $[-3, +3]$ | 4-(b) |
| `training_size` | Percentage of the time-samples in the training set used to train the model | % | 4-(d) |
| HRF | Convolution of the stimuli representation with an Hemodynamic Response Function (HRF) | yes/no | 4-(c) |
| `CNN_layer` | Layer mapped on fMRI data | {`fc7`,<br>`fc7_R`} | 4-(e) |
| `N_iterations` | Number of iterations of the optimiser | 50 – 1000 | 4-(g) |

### 2.3. Decoding with Transfer Learning

By linking deep learning representation with brain data, a straightforward advantage is the possibility to transfer the good discrimination ability of deep networks also to brain data. Once a model has been learned on the training data, we reconstructed the fc7 features of the test images from the fMRI data, and perform on those features classification tasks. In particular, we considered the classification in the movie dataset of the two classes “face” vs “full-body” and the classification of the two classes “person” vs. “no-person” on the images dataset.

To illustrate the effectiveness of transfer learning from CNN to fMRI data, we consider three decoding approaches, learning a model on the training data and evaluating it on the test dataset. We decode categories from a) whole brain fMRI data, b) `fc7` features only, and c) reconstructed deep features (`fĉ7`) obtained from the observed test fMRI data. Please note that in c) we also train on the reconstructed deep features of the training dataset.

The chance level (*i.e.*, the performance obtained when the classifier does not learn any association between data and categories and produces random guesses on the test dataset) can be seen as a “lower bound” for performance, while the decoding in b) can be seen as an “upper bound” as it based on the *true* deep features. We hypothesise that the performance of the decoding analysis using reconstructed deep features (c) will be better than when using imaging data alone (a).

We used a Random Forest (RF) [51] classifier and test the classification performance; RF is used for its capability to deal with big and unbalanced datasets with respect to other methods. For every test we make use of the optimiser described in Section 2.2.1 during the training procedure to select the hyper-parameters (number of trees in the forest, number of used features, and the maximum depth of the tree).

## 3. Results

### 3.1. Linking Methods

The obtained results are shown in Figure 3, which presents the Pearson correlation *r* between `fc7` and `fĉ7` of the image object class “person” on every validation movie, averaged across all features. Since optimising all hyper-parameters (including `time_shift` or the usage of `HRF`) would have led to an explosion of cases, and considering that the choice of some hyper-parameters applies to all linking methods, for this first analysis we hypothesise that `time_shift`= –2 (*i.e.*, `fc7` are delayed of 6*s* with respect to fMRI) as suggested by a previous study which used the same dataset [3]. Other hypotheses are: no use of `HRF`, layer= `fc7_R`, `N_iterations`= 500, and 100% `training size` (where possible). In Section 3.2, for the best method that comes out of this analysis, each of the above hypotheses is tested.

**Figure 3:**
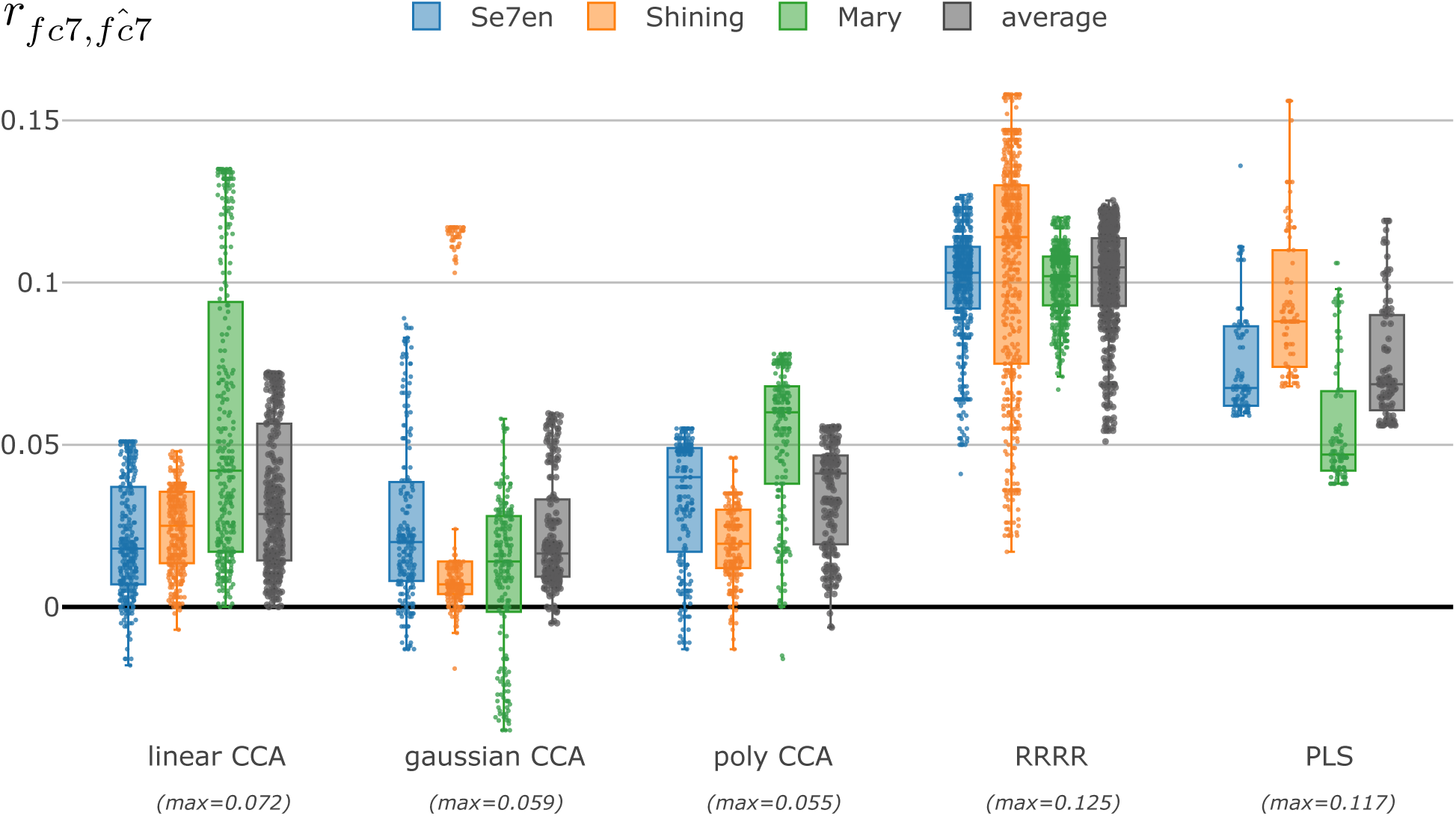
Mapping method comparison in terms of Pearson correlation. Every movie of the validation set (*Se7en, Shining*, and *Mary*) is tested and reported along with the average across movies; every point shows a different step of the optimiser (i.e., a different set of hyper-parameters). Below each name, the maximum value found for every method is reported.

The results show that, while all CCA based methods behave similarly (average Pearson correlation below 0.05), better performance are obtained with PLS and RRRR. In particular, RRRR provides a sensibly better and more stable feature reconstruction across the different validation movies showing an average Pearson correlation larger than 0.1. Therefore, in the remainder of the work, RRRR is chosen as the linking method between CNN features and imaging data.

### 3.2. RRRR hyper-parameter optimisation

The results of a more in-depth optimisation of the RRRR hyper-pameters described in Section 2.2.1 is shown in Figure 4.

**Figure 4:**
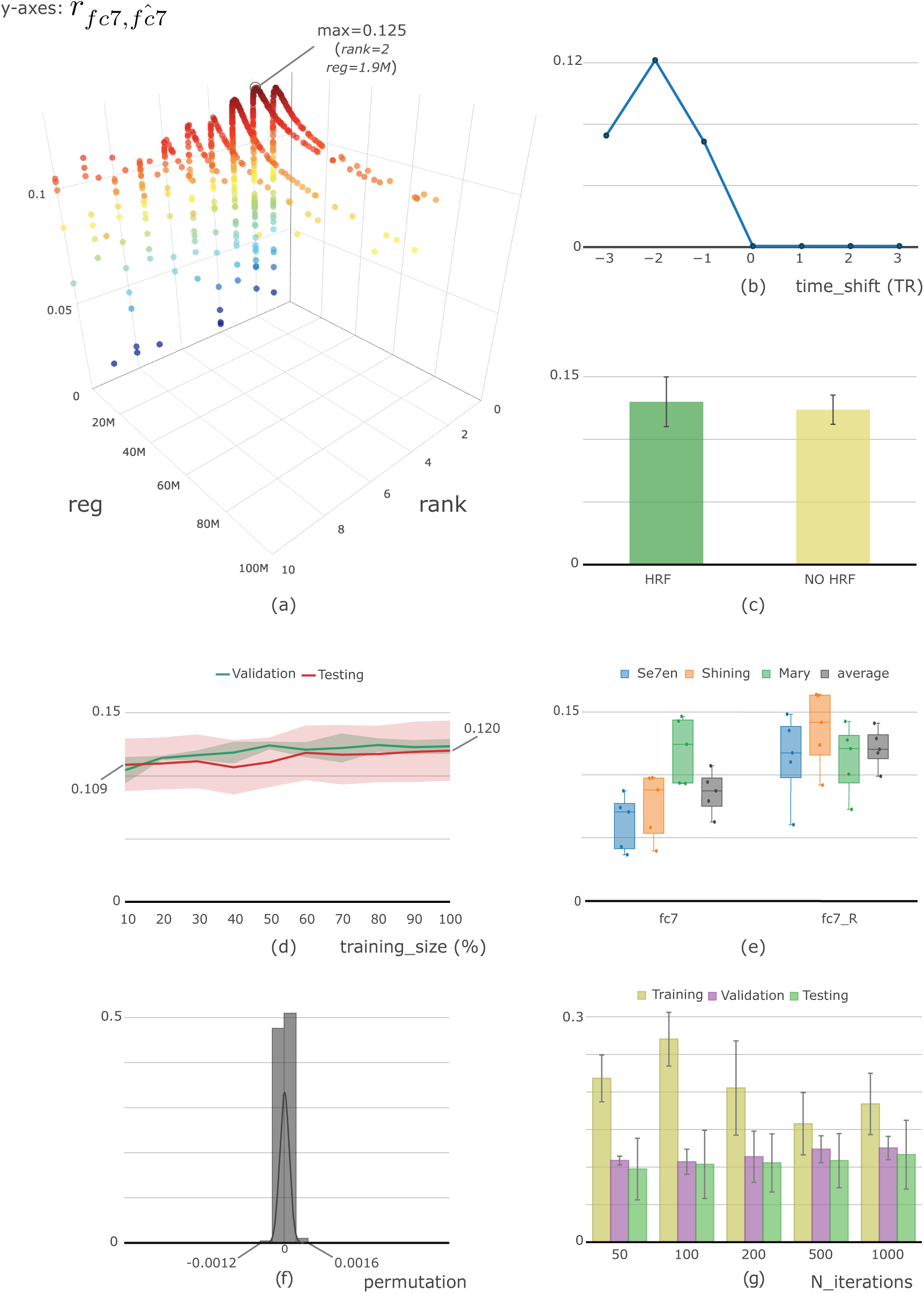
Framework tuning with optimisation of: (a) `rank` and `reg`, (b) `time_shift`, (c) use of hemodynamic response function (`HRF`) or not, (d) training size, (e) `fc7` layer (with and without ReLu), and (g) `N_iterations`. The distribution in (f) depicts the results of the permutation test. All the results are reported in term of Pearson correlation *r*.

#### Rank, reg, and time_shift

The first set of hyper-parameters to be optimised includes rank, `reg` value, and `time_shift`. For every `time_shift` in range [–3, +3] (TR), an optimisation process is launched in order to estimate the other two. Results are shown in Figure 4-(a) and (b), which show different combinations of `rank` and `reg` with the best `time_shift` (= –2), and the best correlation (optimising `rank` and `reg`) found for every `time_shift`, respectively. Value `time_shift` = –2 returns the highest correlation, in line with what is expected from the hemodynamic response, which peaks 4 to 6 seconds after the stimulus onset.

#### HRF

Another decision is whether to use the hemodynamic response function (HRF) or not, which is often convolved with the stimuli representations, in order to ease the mapping. In this case, we compare results with and without convolving `fc7` with HRF (the same used in [3]). To assess the difference, we run two different optimisers in order to find the best correlation value on the validation set changing `rank` and `reg` values (`time_shift`= –2). Results are shown in Figure 4-(c). Despite there is a small improvement in performance by using HRF (mean correlation across movies with HRF = 0.130, without HRF = 0.124) we decide not to continue with this approach. The reason for this is that the visual features could be adversely modified with a convolution with the HRF (that acts as a temporal low-pass filter), potentially reducing the discrimination power of the reconstructed `fc7`, which would not be justified by a marginal increase in correlation.

#### training size

In this work, we are exploiting a very large dataset of fMRI data of subjects watching movies. However, to prove the ability of the method to work well even in (more common) situations in which datasets are smaller, we test our method using different sizes of the training set. Starting randomly selecting only a portion of the training set, from 10% (∼3000 time points) to 100%, we plot the performance in terms of correlation for validation and testing sets (see supplementary material for better details). It is possible to notice two important aspects in Figure 4-(d): both training and testing show well aligned results, proving a very good generality of the method (even with movies and subjects not seen during training), and, in addition, that the performance is good also with relatively small percentages of training set.

#### CNN layer

In CNNs the `fc7` layer is most of the times followed by a rectified linear unit layer (ReLu), an activation function that takes the positive part and thresholds to zero the negative. It is common practise to extract the `fc7` activation before ReLu *i.e.*, with negative values, in those case where we do not have a-priori knowledge about which visual classes there may be in the image. Conversely, as in our case, when a description of a specific image object class is expected, activations are usually taken after ReLu, thus obtaining a sparse representation of the object class. In this work, we select a-priori to use the fully connected layer 7 after ReLu activation function, since we are interested in the image class “person”. However, in Figure 4-(e) we show a comparison between the two approaches, with and without ReLu, noticing that the version without ReLu is doing slightly worse than the counterpart with ReLu (thresholded values). This result is expected, since negative values, thresholded in the case with ReLu, do not carry information about the person class, but force the mapping method to link also these values, thus worsening correlation performance.

#### N_iterations

Finally, to obtain a good training of the model, we need to understand how many iterations our optimiser needs to run in order to reach an optimal solution. The number of iterations is strictly dependent from the search spaces provided to the optimiser: the algorithm needs to know the a-priori probability for every hyper-parameter; the wider the space, the larger number of iterations are needed to converge. In this experiment, we run five different instances with a different number of optimiser iterations, with a common search space of `rank` = *Integer*(1, 100) and `reg` = *Real*(1*e* – 3, 1*e* + 12,“log-uniform”). In Figure 4-(g) training, validation, and testing set performance are shown for 50, 100, 200, 500, and 1, 000 iterations. With such a broad space, a large number of iterations is needed; however, after a certain amount, the improvement is not cost-efficiency positive any more.

#### Permutation

Additionally, to test the robustness of the obtained results, a permutation test is performed: training and validation `fc7` features are randomised by scrambling the phase of their Fourier transform with respect to the original features. The entire training-validation procedure is repeated 3, 000 times on randomly permuted features, and the correlations are calculated. It is worth mentioning that with 3, 000 permutations, the lowest attainable *p*-value, 1/3, 001 (0.0003), is obtained when the correlation values observed in the permutations is never equal or exceeds the correlation obtained on the original data. Figure 4-(f) shows the correlation values obtained with the indication of the maximum (0.0016) and minimum (–0.0012) validation set results found, quite far the other performance shown above.

#### 3.2.1. Reduced rank ridge regression versus feature-wise ridge regression

An interesting comparison, which moves along the analyses of different linking methods shown above, is opposing single {*n* × *m*} regression and {*m*} different {*n* × 1} regressions. These are known in literature as *multivariate regression*, in which multiple independent variables predict multiple dependent variables, in opposition to *multiple regression*, in which multiple independent variables predict one dependent variable. In brain imaging, the multiple regression approach is more frequently employed than the multivariate counterpart, probably for its simplicity.

In this section we show results of this comparison. Adopting the library xgboost [52] for regression, {*m*} (i.e., 4096) different regressions are trained and optimised in terms of the regression value; the optimisation follows the approach described in Section 2.2.1. The search space for the regularisation term is `Real(1e-5, 1e+5)` and 25 iterations are applied to find the best `reg` for every regression. The use of `xgboost` library is motivated by the large number of training to be carry out (4096 × 25) and since the package provides an highly efficient implementation of linear regression.

The obtained results are displayed in Figure 5, where, for every movie in the validation set, correlation between the predicted `fĉ7` and the extracted `fc7` features are shown for all the 4096 regressions grouped together.

**Figure 5:**
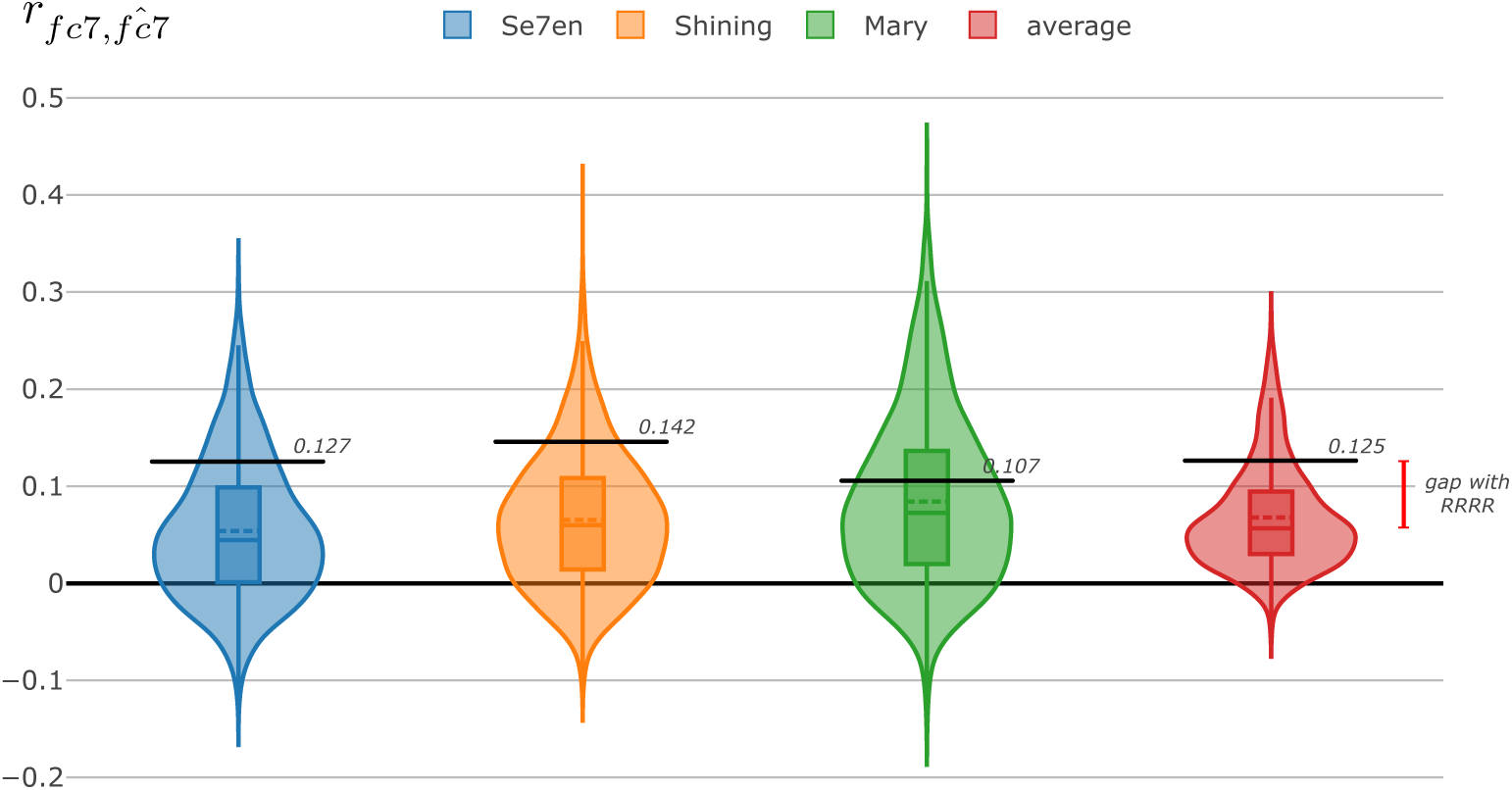
Correlation between `fc7` features and predicted `fĉ7` using optimised multiple ridge regressions. Black lines recall best RRRR results.

Black lines recall RRRR results, highlighting the gap in performance and showing how even if certain {*n* × 1} regressors have high performance (e.g. *r* = 0.4), the mean value of every features predicted is smaller than the mean value obtained for RRRR. A possible explanation of this is that treating each `fc7` separately ignores correlations between them that could aid the prediction.

### 3.3. Correlation on test data

Results are reported for the movie dataset in Figure 6-(a) in terms of average correlation between the predicted `fĉ7` and the extracted `fc7` features, where every dot in the figure is the correlation result for a different subject. The obtained results (*r* = 0.155 as mean correlation for all testing movies, *Poet* = 0.128, *Forrest* = 0.090, *Ryan* = 0.184, *BlackSwan* = 0.063) are remarkable and robust, especially considering that the method is tested across multiple subjects while watching different movie clips not employed during training, and that fMRI data are collected by different MRI scanners.

**Figure 6:**
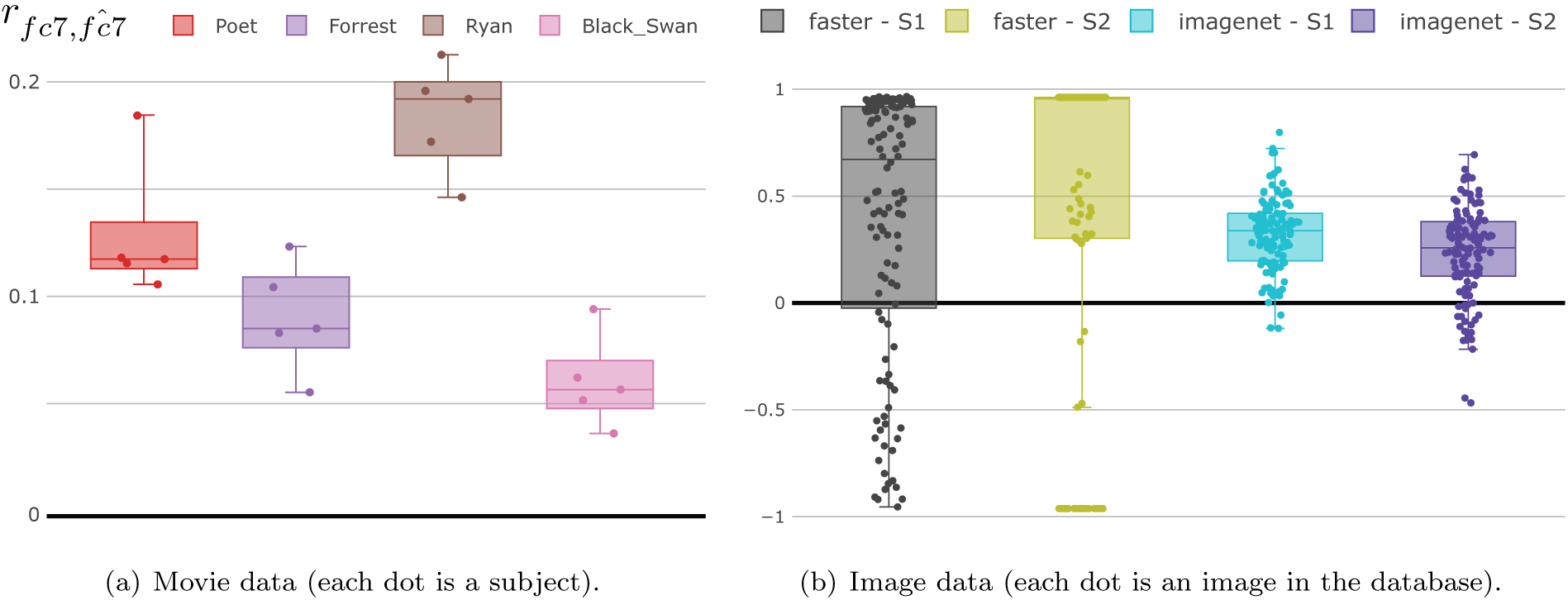
Testing results: (a) correlation results on the testing set (leave out clips) of the movie database (*Poet* = 0.128, *Forrest* = 0.090, *Ryan* = 0.184, *BlackSwan* = 0.063) and (b) correlation on testing images from Kay et al. [11] database (faster r-cnn corr mean: *S*_1_ = 0.390, *S*_2_ = 0.459, vgg-imagenet corr mean: *S*_1_ = 0.323, *S*_2_ = 0.234).

A different test we performed, not shown here for the sake of brevity, switches the roles of the validation and testing sets (*i.e.*, use the testing set of Table 1 as a validation set, and viceversa), to highlight potential differences; however, also in this case, results indicate a mean correlation on (the new) validation set of 0.106 (before it was 0.125), and also the selected values for hyper-parameters are very close to previously obtained ones. The results obtained on the testing set are clearly significant, since the permutation test accomplished on 3, 000 evaluations (and reported in Fig. 4-(f)) never achieved correlation values greater than 0.0016.

The results obtained on the images dataset are shown in Figure 6-(b), where every dot is the correlation result for a particular image. As we described in Section 2.1.2, the database in [11] consists of 1, 750 training and 120 testing images (called “validation” in the original paper) which are provided with the estimated peak BOLD responses (*i.e.*, GLM’s betas). Using the same learning procedure we trained two models to decode two `fc7` activations obtained from two different CNN architectures, namely VGG-16 [38] trained on ImageNet [39], and faster R-CNN [33].

Since brain data available are not registered to any standardised anatomical images, as we have done with movie clips data, the entire training-testing procedure is performed within-subject, and we report results for the two subjects separately (*S*_1_ and *S*_2_). The model is trained relying on the hyper-parameters optimising procedure used before; an optimiser, at every step, measures the performance for a particular set of model hyper-parameters using a 5-fold cross-validation procedure. In particular, due to the within-subject approach, two optimisation procedure instances are carried out, for each of the two subjects, even if similar hyper-parameters are found.

On the left of Figure 6-(b) we show the correlation results obtained by using the same faster R-CNN network used for movie clips (corr mean: *S*1 = 0.390, *S*2 = 0.459), while on the right of the same figure we show performance obtained with the VGG-16 network trained on ImageNet (corr mean: *S*1 = 0.323, *S*2 = 0.234). We chose to test two different `fc7` features because in the case of faster R-CNN network we wanted to perform further classification on the object class “person” as done with movies. Conversely, by extracting features by a VGG network trained on ImageNet we aimed at measuring correlation on `fc7` features potentially descriptive for any type of object class among those present in ImageNet database.

### 3.4. Decoding with Transfer Learning

The results of the decoding analyses are reported in terms of *balanced accuracy*. With respect to the commonly used accuracy (i.e., the number of correct predictions divided by the total number of tested samples) balanced accuracy is computed as the average of single class accuracy, and has been advocated as a better performance metric when there is a strong unbalance between class cardinalities [53, 54].

Results on movies and reported in Figure 7-(a) indicate how using `fĉ7` is positioned between `fc7` and fMRI data. Among these bars, the key comparison is the one between the classifier using fMRI data only (testing set overall mean = 51.1%) and the `fĉ7` based classifier (mean = 59.6%), showing a relevant difference and good generalisability across subjects. Results on fĉ7 are quite close to those obtained with the original `fc7` (overall mean = 65.1%). Figure 7-(b) shows the balanced accuracy results for tested images of the images dataset for the two subjects S1 and S2: in this case, the difference is not remarkable as it as happens with movies, and the classification using predicted `fĉ7` features works well with only one subject. This may mean that the poor correlations results (negative values reported for S2 in Fig. 6) have a negative contribution in the classification process. Classification results in terms of accuracy, balanced accuracy, and confusion matrix, for movie and image data, are full reported in supplementary material.

**Figure 7:**
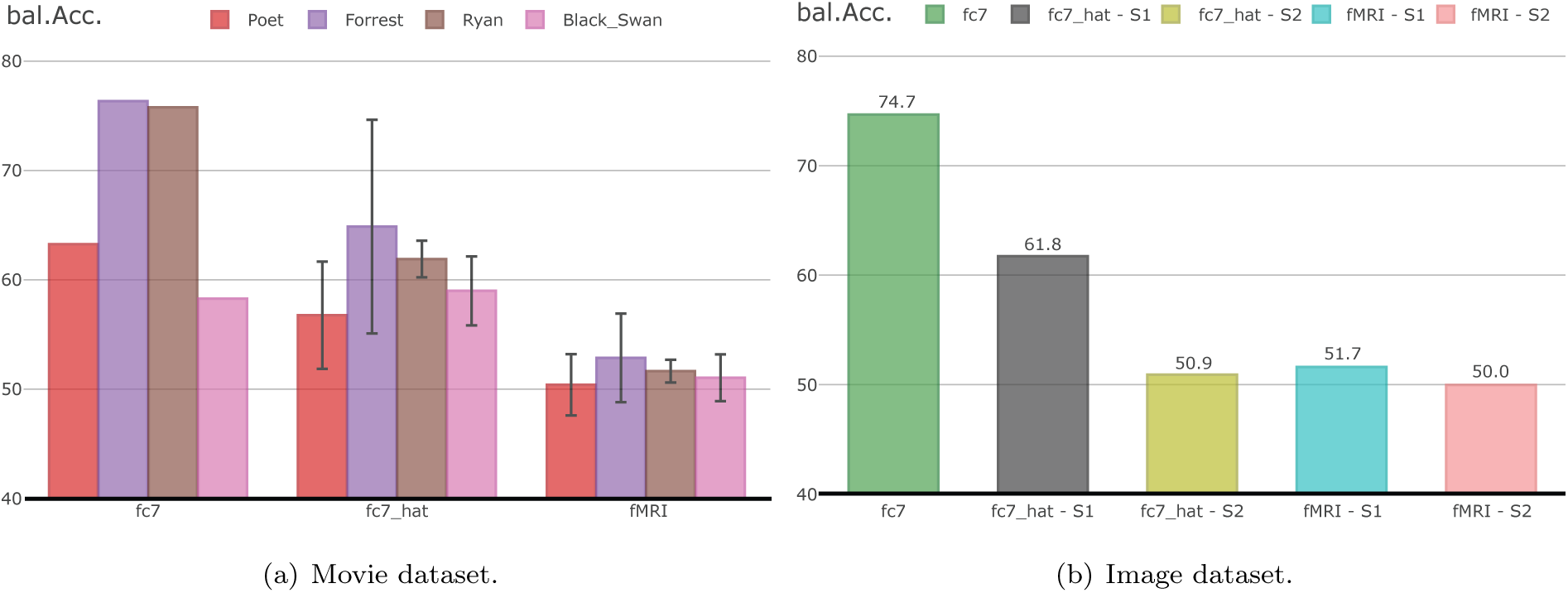
Classification results, in terms of balanced accuracy, on: (a) “face” vs “full-body” on testing movie clips, and (b) “person” vs “no-person” on the testing set from Kay at el. [11].

### 3.5. Imaging subspace projection

The reduced `rank` ridge regression (RRRR) approach is based on the projection of high dimensional imaging data onto a subspace of lower dimension. Since this projection is linear, it is possible to visualise the projections that are applied to imaging data to reconstruct the deep features (see Appendix A for more details). As the hyperparameter tuning suggested optimal results with `rank` = 2 (see Figure 4), we display in Figure 8 two projection maps, one per dimension of the reduced space (see also Table 9-10 in Supplementary Material).

**Figure 8:**
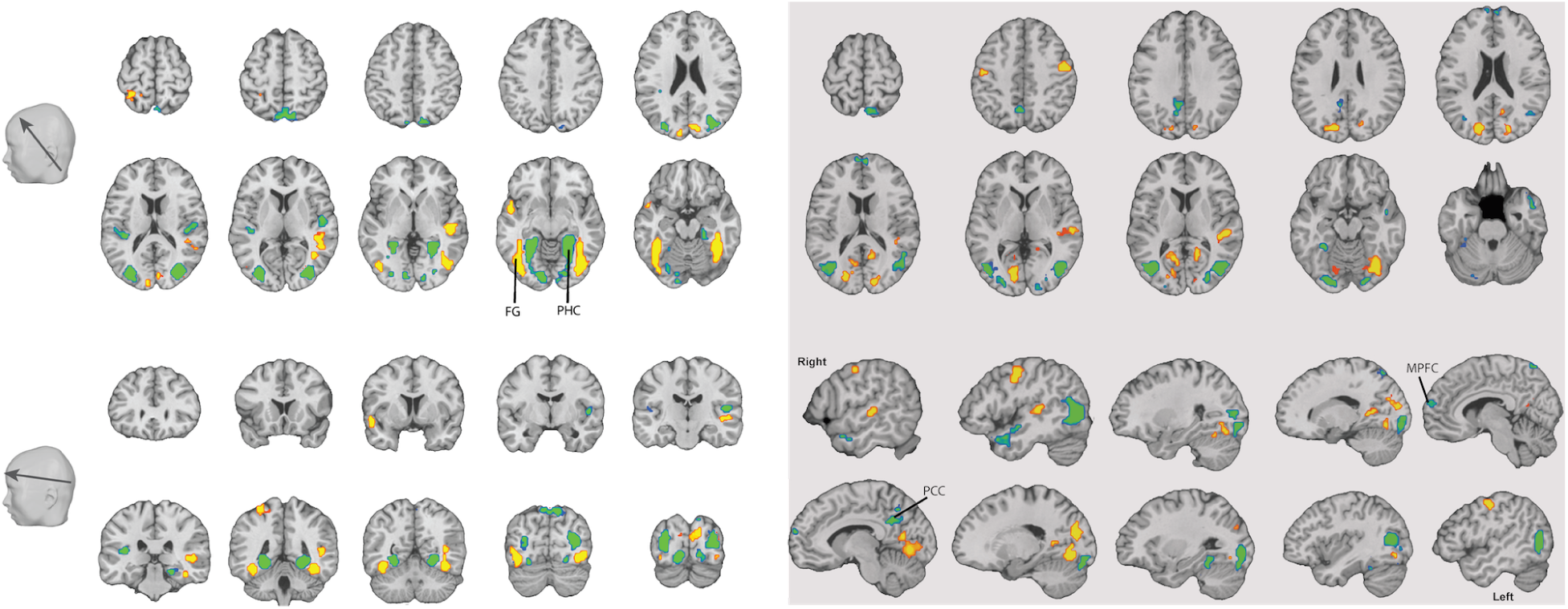
Three dimensional maps of the RRRR projections. Only the top 5% of the weights are visualised (minimal cluster size: 25 voxels). Abbreviations: FG - fusiform gyrus; MPFC - medial prefrontal cortex; PHC - parahippocampal cortex; PCC - posterior cingulated cortex.

For the sake of clarity, we present the top 5% of the weights in these models. The first projection (left panel) includes major bilateral clusters with opposite signs in the fusiform gyrus (including the fusiform face area) and the parahippocampal pyrus (including the parahippocampal place area). These regions have been associated with the face processing [55] and scene recognition [56], respectively. The second projection (right panle) included major hubs in the motor cortex (bilateral) and association visual (bilateral) and auditory (right) cortex. It also included large clusters across the posterior and anerior superior temporal cortex, medial prefrontal cortex and the posterior cingulated cortex, which have been implicated in social cognition and mentalization [57].

To assess the functional meaning of the model in a quantitative manner, we used the web-based multi-study decoder NeuroVault [58], which allows for the interpretation whole-brain patterns based on a large database of neuroimaging studies. The top functional entries that were associated by the decoder with the first projection were “face”, “recognition”, and “face recognition”. The second projection was most strongly associated with the entries “vocal”, “production”, “saccades”, and “speech production”. These findings support the notion that the models captured relevant features in the movies; namely, face presence in the case of the first component and human speech in the case of the second component.

## 4. Discussion and future directions

In this work we have shown how to harness the richness of deep learning representations in neuroimaging decoding studies. The potential benefit of the use of CNNs derived features *i.e.*, `fc7`, is twofold. First, it is possible to perform a task *i.e.*, the regression from imaging data to `fc7`, that is more manageable, from the dimensionality point of view, than a simple classification based solely on imaging data. We have shown in Section 2.3 that this approach is useful and results in better classification performance, demonstrating how to embed high-dimensional neuroimaging data onto a space designed for visual object discrimination. In addition, using these networks for feature extraction allows us to extract stimuli representations by means of an automatic procedure that does not require ground truth or supervision, and that may help to more easily address certain unsupervised learning tasks.

Regarding the good classification performance achieved with the predicted `fĉ7`, it is possible that this is due to intrinsic redundancy and sparsity properties of CNN representations. A good analogy may be the signal transmission process, in which some redundancies are introduced on purpose before transmitting the information through the channel, so that the overall process can afford some losses. Also in this case the CNN redundancy allows to obtain good classification performance despite the fact that the reconstruction `fĉ7` is not perfect.

In brain imaging literature, and in a broader sense in all biomedical engineering fields, from neuroscience to genetic, there are plenty of multivariate linking methods, with different formulations and training strategies. In this work we have compared some of the most widely used multivariate approaches, and our results indicate that the best performance for the type of data we considered is obtained by RRRR method. Given the large amount of time points on which it has been tested, these results are reliable and we thus recommend the use of RRRR in the context of combining fMRI data and such deep computational models.

The reliability of the proposed method is clearly demonstrated by the adopted inter-subject approach in the movie dataset. While most of the works present in literature rely on single subject analyses in very controlled stimulation settings, we decided to also consider, alongside with the static image dataset, movie clips with free viewing. In addition to this, we performed training and testing using separate movie subsets, testing new subjects on unseen stimuli, and employing data coming from different MRI scanners. The good results shown above can be seen as a positive assessment of the across-subject registration process to standardised anatomical images (done with Brainvoyager QX, Brain Innovations). This because the conducted analyses give us a quantitatively remarkable confidence about the fact that the functional responses are aligned, which means that the anatomical inter-subject alignment succeeds at the spatial scale suited for performing the considered decoding task. However, more fine grained discrimination may require different, and more advanced, alignment procedures that take into account anatomical differences and anatomical/functional misalignment across subjects [59].

The free viewing condition (no fixation imposed during the experiment), can be seen as a shortcoming of the work on the movie dataset, since there is no direct correspondence between cortical space in early vision cortex and the image space. However, within a time resolution of 3 s., we can be quite confident that the participants watched the same areas in the movie frames. Apart from few people-less movies, such as documentaries, the presence of human figures is central to modern cinematography [60, 61]. Different studies investigated fixation similarities of subjects while watching dynamic scenes and, most of them, led to the same conclusion: that movies, especially Hollywood clips, are able to intrinsic catch the attention of the viewers on the same parts of the movie frame [34, 35, 62, 63]. This is specially true if there is a temporal “average” of a few seconds as it happens in fMRI. If, on the one hand, this type of stimulus can give us high confidence to the fact the viewer attention will be focused on “person” class (and related ones, such as “face”, “human-body”, and “no-person”) in a free-viewing experiment, on the other hand we have to face up the strong limitation regarding other object classes, since we cannot be sure that the viewer attention would focus on different scene objects. This, however, is a limitation of the dataset we employed, whereas one could envision experiments where stimuli of categories different from faces are considered proposed approach

In this work we investigated the reconstruction of CNN features from imaging data for the purpose of decoding visual stimulus categories (thus performing *decoding*). However, a natural extension of the work would be to reconstruct imaging data based on CNN features (*i.e.*, inverting the directionality of the Reduced Rank Ridge Regression used); this would be similar to current encoding approaches, with the difference of performing a multivariate regression, as compared to massive univariate regressions traditionally used, which could be better suited in handling correlation structures in the feature space. The comparison between RRRR and Ridge Regression suggests that this could hold true also when performing encoding, but further work is needed to test this explicitly.

## 5. Conclusion

In this work we proposed a robust method for decoding DL features from brain imaging data (fMRI). Whereas the direct application of CNN architectures to decipher subject states or perception from imaging data is dramatically limited by the relative scarcity of available brain data, it is still possible to improve fMRI-based decoding benefiting from the non-linear feature mapping of CNNs by means of transfer learning.

We have shown how to establish a multivariate link between the imaging data (fMRI) and the first fully connected layer (`fc7`) of a CNN, which enables deep feature decoding. To this end, we use Reduced Rank Regression with Ridge Regularization (RRRR), that is particularly suitable in handling high dimensional, correlated, voxels time series and `fc7` features. We validated and exploited the `fc7` decoded features, performing an object image classification task on two classes (*i.e.*, “face” vs “full-body”, and “person” vs “no-person”) on two different databases, one based on static images and the other based on a large cohort of movie based scans. We compared the obtained classification with other methods using shallow machine learning classifiers that do not exploit the richness of the deep representation. Results confirmed the reliability of the established mapping between fMRI data and CNN layers to provide good representations of visual stimuli, which can be used as a generic mapping method for further research in visual decoding and encoding.

## Appendix A. RRRR formulation

The formulation and estimation of the Reduced Rank Ridge Regression used in this work is provided in [47]. We summarise here the aspects most relevant for the current work.

Denote as *X* the [*n* × *p*] fMRI data, where *n* is the number of time points and *p* the number of voxels. The deep features are represented in *Y*, of size [*n* × *q*], where *q* denotes the number of features. The goal of Reduced `Rank` Ridge Regression is solve the problem

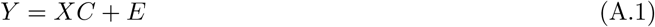

where *C* is [*p* × *q*] contains the regression coefficients and *E* is the [*n* × *q*] error matrix.

The solution proposed in [47], based on subspace projection on a space of lower dimension (*reduced rank*) togher with L2 regularisation (*ridge*) is:

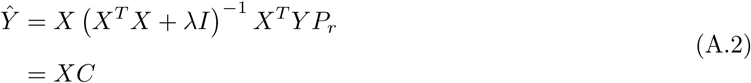

where *P*_*r*_ is the projection matrix, i.e., the matrix that projects the matrix *Y* to an *r*-dimensional space. The coefficient matrix *C* (of rank *r* ≤ min(*p, q*)) can be decomposed using an “economical” version of Singular Value Decomposition (SVD) (where all zero eigenvalues are removed from the decomposition) as a product of two matrices of rank *r*:

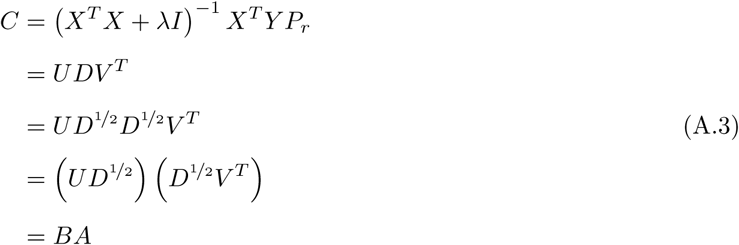

obtaining two terms *A* and *B* of dimensions respectively of [*r* × *q*] and [*p* × *r*]. Please note that the decomposition of *D* = *D*^½^ *D*^½^ is valid since *D* is a diagonal and square matrix.

The regression in (A.1) can now be interpreted in the following way:

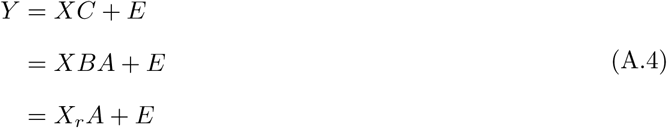

where *X*_*r*_ is a [*n* × *r*] matrix containing the *projection* of fMRI data onto a space of dimension *r*. These components are then combined using matrix *A* to reconstruct the deep features in *Y*. The maps visualised in Section 3.5 show the mapping performed by *B*.

## Footnotes

1 Code: http://github.com/rockNroll87q/RRRR

