## supplementary for "Transfer learning of deep neural network representations for fMRI decoding"

### 1. Training size

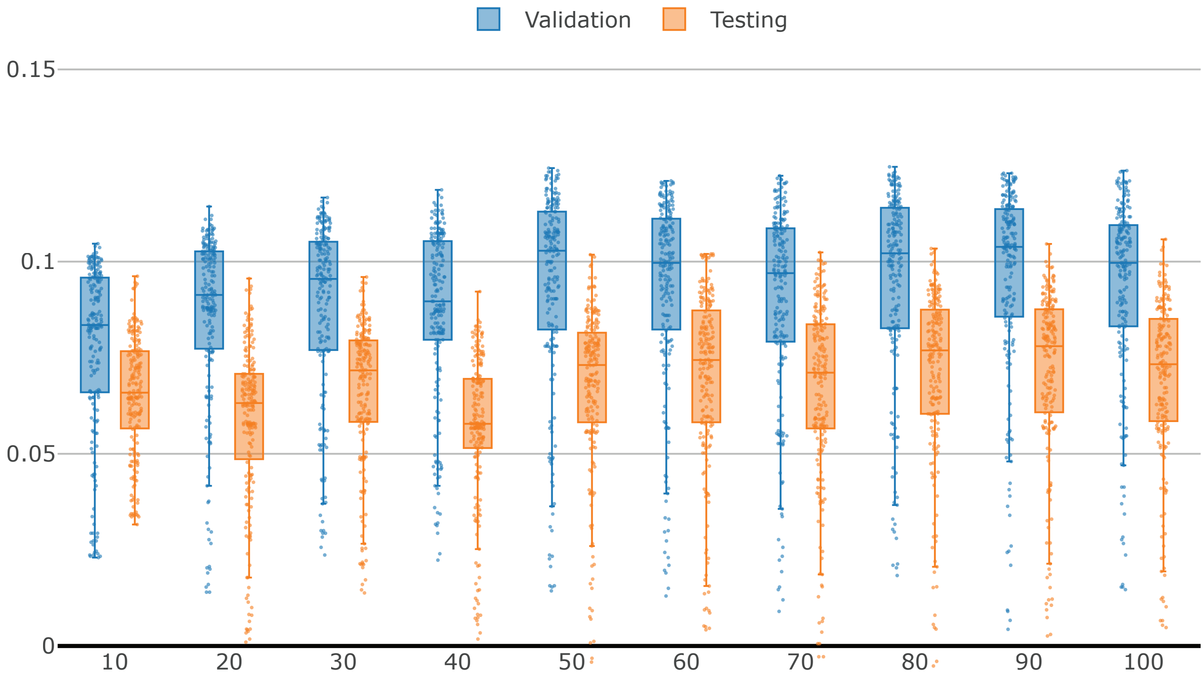


Figure 1: Framework tuning with optimisation of `training_size`. Results on validation and testing dataset are shown. Maximum values for every column are reported in Figure 3-(c) of the paper.

*Corresponding author
*Email address:* `Michele.Svanera at glasgow.ac.uk` (Michele Svanera)

### 2. Classification results

#### *2.1. Movie data*

Using `fc7`: accuracy = 71.11%, balanced accuracy = 65.07%.

Table 1: Classification on movie data. Confusion matrix on Test-set using `fc7`.

| | **"face"** | **"full-body"** |
|---|---|---|
| **"face"** | 428 | 414 |
| **"full-body"** | 431 | 1652 |

Using $\hat{\texttt{fc7}}$: accuracy = 64.92%, balanced accuracy = 59.62%.

Table 2: Classification on movie data. Confusion matrix on Test-set using $\hat{\texttt{fc7}}$.

| | **"face"** | **"full-body"** |
|---|---|---|
| **"face"** | 397 | 445 |
| **"full-body"** | 581 | 1502 |

Using VTC: accuracy = 70.01%, balanced accuracy = 51.07%.

Table 3: Classification on movie data. Confusion matrix on Test-set using VTC.

| | **"face"** | **"full-body"** |
|---|---|---|
| **"face"** | 54 | 788 |
| **"full-body"** | 89 | 1994 |

#### *2.2. Image data*

Using `fc7`: accuracy = 89.17%, balanced accuracy = 74.68%.

Table 4: Classification on movie data. Confusion matrix on Test-set using `fc7`.

| | **"person"** | **"no-person"** |
|---|---|---|
| **"person"** | 96 | 3 |
| **"no-person"** | 10 | 11 |

**S1**. Using $\hat{\texttt{fc7}}$: accuracy = 61.67%, balanced accuracy = 61.76%.

Table 5: Classification on movie data. Confusion matrix on Test-set using $\hat{\texttt{fc7}}$.

| | **"person"** | **"no-person"** |
|---|---|---|
| **"person"** | 61 | 38 |
| **"no-person"** | 8 | 13 |

**S2**. Using $\hat{\texttt{fc7}}$: accuracy = 71.67%, balanced accuracy = 50.94%.

Table 6: Classification on movie data. Confusion matrix on Test-set using $\hat{\texttt{fc7}}$.

| | **"person"** | **"no-person"** |
|---|---|---|
| **"person"** | 82 | 17 |
| **"no-person"** | 17 | 4 |

**S1**. Using VTC: accuracy = 45.00%, balanced accuracy = 51.65%.

Table 7: Classification on movie data. Confusion matrix on Test-set using VTC.

| | **"person"** | **"no-person"** |
|---|---|---|
| **"person"** | 41 | 58 |
| **"no-person"** | 8 | 13 |

**S2**. Using VTC: accuracy = 17.50%, balanced accuracy = 50.00%.

Table 8: Classification on movie data. Confusion matrix on Test-set using VTC.

| | **"person"** | **"no-person"** |
|---|---|---|
| **"person"** | 0 | 99 |
| **"no-person"** | 0 | 21 |

Table 9: Size and Talairach coordinates of the **First Column** employed for decoding the movie's features.

| **Region label** | **Cluster size** | **X** | **Y** | **Z** |
|---|---|---|---|---|
| Positive weights | | | | |
| R Fusiform Gyrus, Inferior Occipital Gyrus | 314 | 39 | -46 | -17 |
| L Fusiform Gyrus, Inferior Occipital Gyrus | 254 | -42 | -79 | -11 |
| R Superior Temporal Gyrus | 99 | 45 | -25 | -2 |
| L Superior Temporal Gyrus | 36 | -54 | 5 | -8 |
| R Superior Temporal Gyrus | 22 | 48 | 2 | -14 |
| L Postcentral Gyrus | 29 | -30 | -43 | 64 |
| R Cuneus | 37 | 12 | -85 | 22 |
| L Cuneus | 25 | -9 | -91 | 19 |
| Negative weights | | | | |
| R Parahippocampal Gyrus | 264 | 27 | -46 | -5 |
| L Parahippocampal Gyrus | 273 | -27 | -49 | -5 |
| R Middle Occipital Gyrus | 216 | 33 | -79 | 13 |
| L Middle Occipital Gyrus | 166 | -33 | -88 | 10 |
| R Transverse Temporal Gyrus | 61 | 54 | -16 | 10 |
| L Transverse Temporal Gyrus | 49 | -36 | -34 | 13 |
| L Cingulate Gyrus | 21 | -9 | -22 | 37 |
| R+L Precuneus | 116 | 15 | -79 | 49 |
| L Inferior Frontal Gyrus | 21 | -36 | 26 | -5 |

Table 10: Size and Talairach coordinates of the **Second Column** employed for decoding the movie's features.

| **Region label** | **Cluster size** | **X** | **Y** | **Z** |
|---|---|---|---|---|
| Positive weights | | | | |
| R Lingual Gyrus | 109 | 33 | -70 | -17 |
| L Lingual Gyrus | 297 | -15 | -76 | -8 |
| R Precentral Gyrus | 59 | 54 | -7 | 46 |
| R Posterior Cingulate | 42 | 15 | -55 | 10 |
| R Superior Temporal Gyrus | 64 | 48 | -28 | 7 |
| L Precentral Gyrus | 39 | -45 | -13 | 49 |
| R Middle Occipital Gyrus | 43 | 15 | -88 | 16 |
| Negative weights | | | | |
| L Middle Occipital Gyrus, Inferior Occipital Gyrus | 310 | -45 | -76 | 4 |
| R Inferior Temporal Gyrus, Superior Temporal Gyrus, Inferior Occipital Gyrus, Middle Occipital Gyrus | 339 | 48 | -76 | 1 |
| R Middle Temporal Gyrus | 54 | 51 | 5 | -26 |
| L Precuneus | 58 | -12 | -49 | 28 |
| L Cerebellum (Culmen) | 52 | -33 | -43 | -23 |
| R+L Dorsomedial prefrontal cortex | 26 | 6 | 62 | 19 |
| R Superior Parietal Lobule | 34 | 6 | -67 | 61 |
